# Vesicle docking and fusion pore modulation by the neuronal calcium sensor Synaptotagmin-1

**DOI:** 10.1101/2024.09.12.612660

**Authors:** Maria Tsemperouli, Sudheer Kumar Cheppali, Felix Rivera Molina, David Chetrit, Ane Landajuela, Derek Toomre, Erdem Karatekin

**Author notes:** These authors contributed equally to this work (Maria Tsemperouli, Sudheer Kumar Cheppali). Present address: Solar Energy for Net Zero Research Cluster, School of Engineering, University of British Columbia, Kelowna, BC Canada. **CONFLICT OF INTERESTS:** The authors declare no competing financial interests.

## Abstract

Synaptotagmin-1 (Syt1) is a major calcium sensor for rapid neurotransmitter release in neurons and hormone release in many neuroendocrine cells. It possesses two tandem cytosolic C2 domains that bind calcium, negatively charged phospholipids, and the neuronal SNARE complex. Calcium binding to Syt1 triggers exocytosis, but how this occurs is not well understood. Syt1 has additional roles in docking dense core vesicles (DCV) and synaptic vesicles (SV) to the plasma membrane (PM) and in regulating fusion pore dynamics. Thus, Syt1 perturbations could affect release through vesicle docking, fusion triggering, fusion pore regulation, or a combination of these. Here, using a human neuroendocrine cell line, we show that neutralization of highly conserved polybasic patches in either C2 domain of Syt1 impairs both DCV docking and efficient release of serotonin from DCVs. Interestingly, the same mutations resulted in larger fusion pores and faster release of serotonin during individual fusion events. Thus, Syt1’s roles in vesicle docking, fusion triggering, and fusion pore control may be functionally related.

## INTRODUCTION

Synaptotagmin-1 (Syt1) is a major calcium sensor that promotes rapid neurotransmitter release in neurons and hormone release in many neuroendocrine cells^1–3^. It has two cytosolic C2 domains that bind calcium, negatively charged phospholipids, and the neuronal SNARE complex (Figure 1A). Calcium binding to Syt1 triggers exocytosis, but how this occurs is poorly understood^1–8^.

**Figure 1.**
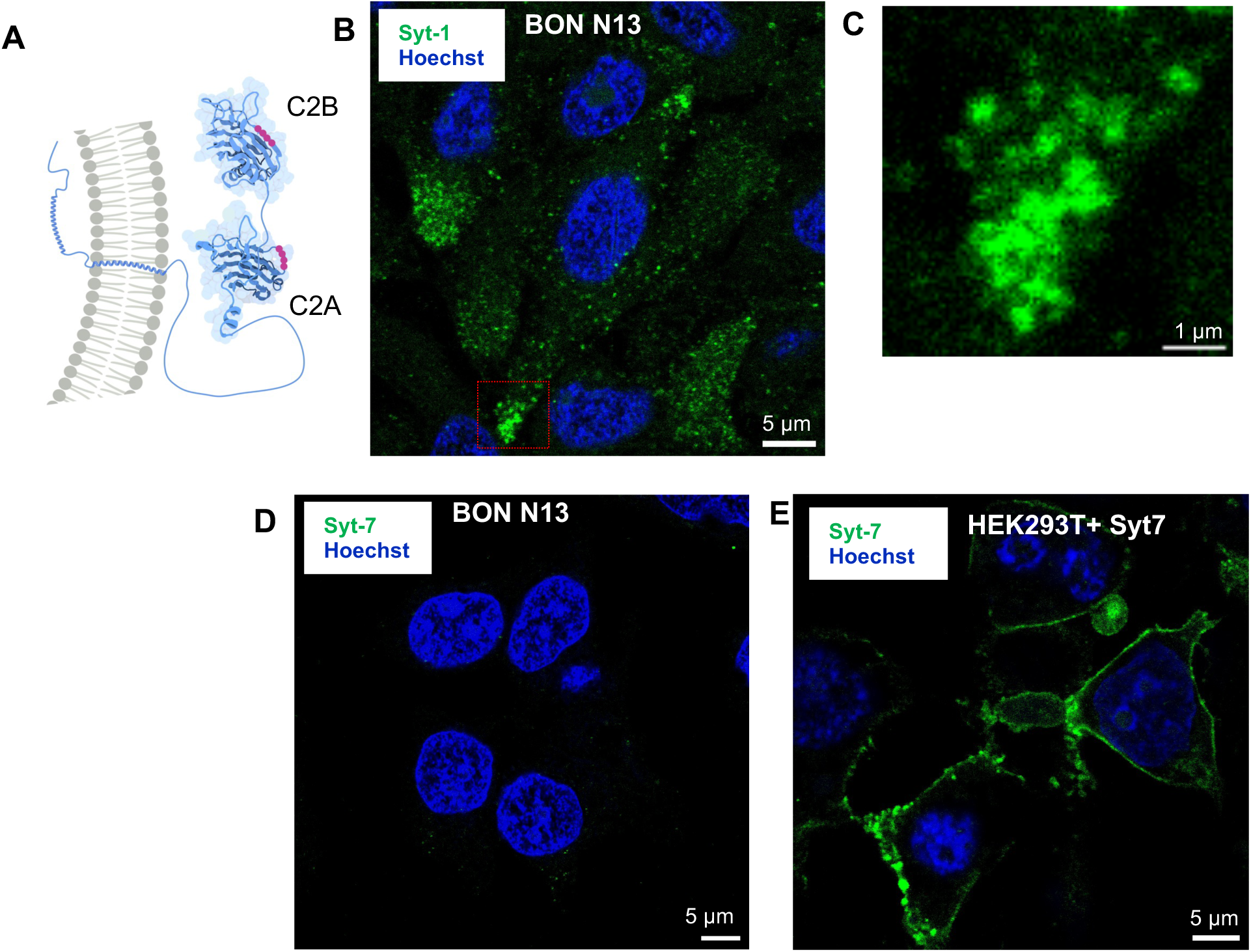
Synaptotagmin-1, but not Synaptotagmin-7, is expressed in BON cells. **A.** Schematic structure of Syt1 with its polybasic patches, marked as purple discs. The C2A and C2B domains are rendered using bioRender (bioRender.com) from the Protein Data Bank entries, 3F04 and 1K5W, respectively. The rest of the molecule is schematically drawn using bioRender (bioRender.com). **B**. Immunofluorescence of BON cells using anti-Syt1 antibodies. Cell nuclei were labeled with Hoechst dye (blue). Syt1 immunofluorescence (green) appears punctate, consistent with the distribution of DCVs. **C**. An enlarged view of the red boxed region in B, showing a cluster of puncta. **D**. No immunofluorescence against Syt7 could be detected in BON cells. **E**. As a positive control, HEK293T cells transiently expressing Syt7 displayed robust Syt7 immunofluorescence.

Most prior research focused on the role of Syt1 in rapid calcium-triggering of release. But Syt1 has also been implicated in docking of dense core vesicles (DCV) and synaptic vesicles (SV), although some reports are inconsistent for the latter. Bommert et al.^9^ reported that injection of blocking peptides increased the number of SVs within 50 nm of the plasma membrane (PM) in the squid giant presynaptic terminal. In the *C. elegans* neuromuscular junction (NMJ), Syt1 KO results in decreased SV density^10^ and docking defects^11^. In the Drosophila NMJ, Syt1 KO leads to a reduction in the number of docked SVs^12^. In hippocampal neuronal mass cultures from Syt1 KO mice, Geppert et al.^13^ did not detect any qualitative changes in SV density or docking compared to cultures from wild-type mice. By contrast, Liu et al reported both reduced SV density and docking for Syt1 KOs in a similar preparation^14^. Analyzing organotypic mass cultures of hippocampal neurons, Imig et al. concluded that Syt1 KO does not have any major direct effects on SV docking^15^. Yet, Chang et al.^16^, found that a polybasic patch in the C2B domain of Syt1 (K325-327) plays a role in SV docking in cultured hippocampal neurons.

Similarly, Chen et al.^17^ reported that C2B polybasic patches are required for SV docking in hippocampal neuronal cultures and suggested PI(4,5)P_2_ is the docking partner on the PM, not the t-SNAREs Syntaxin-1 and/or SNAP25. In contrast to these reports on SV docking, DCV docking studies indicate an unambiguous requirement for Syt1, likely because vesicle docking phenotypes are much stronger in neuroendocrine cells^18–22^. In adrenal chromaffin cells from Syt1 KO mice, De Wit et al.^20^ found very pronounced DCV docking deficiencies and reported that docking required Syt1 to interact with SNAP25. Corroborating these findings, Mohrmann et al.^23^ further showed that a group of acidic amino acids near the center of the SNARE domains in SNAP-25A is essential for interactions with Syt1 and DCV docking. Syt1 residues responsible for binding SNAP-25A likely involve the C2 domain polybasic patch^24^, but these residues also bind acidic lipids^25–31^. In summary, Syt1 is required for DCV docking, and likely for SV docking as well, but there is uncertainty about the PM docking partner, with both PI(4,5)P_2_ and t-SNAREs being implicated.

Why is Syt1’s role in vesicle docking important? If Syt1 plays a role in vesicle docking, then some of its previously reported effects on rapid exocytosis may need to be revisited: Syt1 could increase the tightly docked pool of vesicles or mediate the rapid release from this pool, or both. This has wide-ranging implications for short-term plasticity^32–36^.

Syt1 additionally plays an important role in regulating fusion pore dynamics^37–43^. The fusion pore is the initial 1-3 nm pore that forms after membrane merger and is a dynamic structure that can flicker open-closed, expand, or reseal^44–47^. Fusion pore dynamics determine release kinetics, the amount of cargo released, and in neuroendocrine cells, the cargo size that can be released^44–47^. Using a nanodisc (ND)-cell fusion assay, we showed that Syt1 promotes fusion pore expansion in cooperation with SNARE complexes and PI(4,5)P_2_, *via* a calcium-dependent lever action that increases the distance between the fusing membranes^37^. This incurs a bending energy penalty, relieved by expansion of the pore diameter^37^. But how Syt1 regulates fusion pore dynamics in live cells remains to be established.

In cultured mouse cortical neurons, we previously found that charge neutralization mutations in the polybasic patches of Syt1 in the C2A (Sty1^K189–192A^) and C2B domains (Syt1^K326,327A^) impaired evoked release^28^. Neither spontaneous release nor the readily releasable pool (RRP, probed by hypertonic sucrose) were significantly affected, suggesting the mutations lowered the probability of release per action potential, P_*r*_. In addition, we had detected a small but significant delay in release kinetics following depolarization^28^. Given that the C2B polybasic patch was previously implicated in SV docking^16,17^, and that the C2A domain bears a similar, highly conserved polybasic patch^28^, we hypothesized that both the C2A and C2B polybasic patches may be important for vesicle docking and that a docking defect might explain the slight delay we observed in release upon stimulation^28^. However, SV docking defects are challenging to detect, as SVs are tightly packed and tethered to one another in addition to the presynaptic PM^36,48^. These technical challenges may explain, at least partially, the inconsistent results regarding the role of Syt1 in SV docking mentioned above. By contrast, DCV docking defects are much easier to detect, due to the lower DCV densities, and perhaps the cortical actin dynamics that move weakly attached DCVs away from the PM^19,22,49,50^.

We thus decided to probe the roles of both the C2A and C2B polybasic patches in DCV docking in neuroendocrine cells. This additionally allowed us to study the dynamics of single fusion pores with high time resolution. Using amperometry and electron microscopy, we found that highly conserved polybasic patches on both the C2A and C2B domains of Syt1 contribute to both vesicle docking and fusion pore regulation.

## RESULTS

### Syt1 is the major calcium sensor for human neuroendocrine BON cell exocytosis

We used human neuroendocrine BON cells^51–53^, which store and secrete, in a calcium-dependent manner DCV contents that includes: serotonin (5-hydroxytryptamine, 5-HT), chromogranin A, neurotensin, and other regulatory peptides and compounds^53,54^. Secretion can be induced by acetylcholine^53–55^, phorbol esters or forskolin^56,57^, the calcium ionophore ionomycin^56–61^, permeabilization in the presence of a calcium buffer^62,63^, or calcium uncaging^62^, but response to depolarization is weak^54^. BON cells have been used as a model cell for studying mechanisms of calcium-regulated exocytosis^49,54,59–64^, as they possess the canonical exocytic machinery, including Munc-18, Munc-13, CAPS, Complexin-1, Synaptotagmin-1, and the SNARE proteins Syntaxin-1, SNAP-25, and VAMP-1^61^. The large size of their DCVs (∼300 nm diameter^62^) facilitate imaging^19,22,49,54,62–64^ and electrochemical detection of release^59,60,64^. Unlike PC12 cells (a widely used model), they have no small synaptic-like vesicles, which can confound interpretation.

We first established that Syt1 is a major calcium sensor in BON cells (Fig 1). Immunolabeling Syt1 produced robust punctate signals (Fig. 1B,C), indicating abundant expression on DCVs. A challenge with adrenal chromaffin cells is that they have distinct DCV populations bearing Syt1 or Syt7, with little overlap^43^; Syt1 and Syt7 have distinct calcium affinities and confer different fusion properties to single DCVs^43^. This dual DCV population greatly complicates the analysis and interpretation of secretion studies in which the two populations cannot be directly distinguished, such as with electrical detection of secretion. We therefore tested for the presence and distribution of Syt7 in BON cells. Syt7 immunolabeling did not produce any detectable signals in BON cells (Fig 1D). As a positive control, we used HEK292T cells transiently expressing Syt7, which produced strong immunofluorescence against Syt7 (Fig 1E). Thus, we attribute the lack of signals in BON cells to the low abundance of Syt7 in these cells.

To determine the role of Syt1 in BON cells DCVs secretion, we then generated BON cell lines in which *syt1* expression is constitutively knocked down (KD) by stable *syt1* depletion with shRNA sequence against it (see Materials and Methods). The estimated *syt1* KD efficiency on the stable shRNA cells was ∼50% after Western blot analysis (Fig 2B,C). Bulk release of serotonin, induced by ionomycin, and measured by ELISA, was impaired to nearly the same extent (Fig. 1D), suggesting that Syt1 is a major calcium sensor for exocytosis in BON cells, consistent with a previous screen^61^.

**Figure 2.**
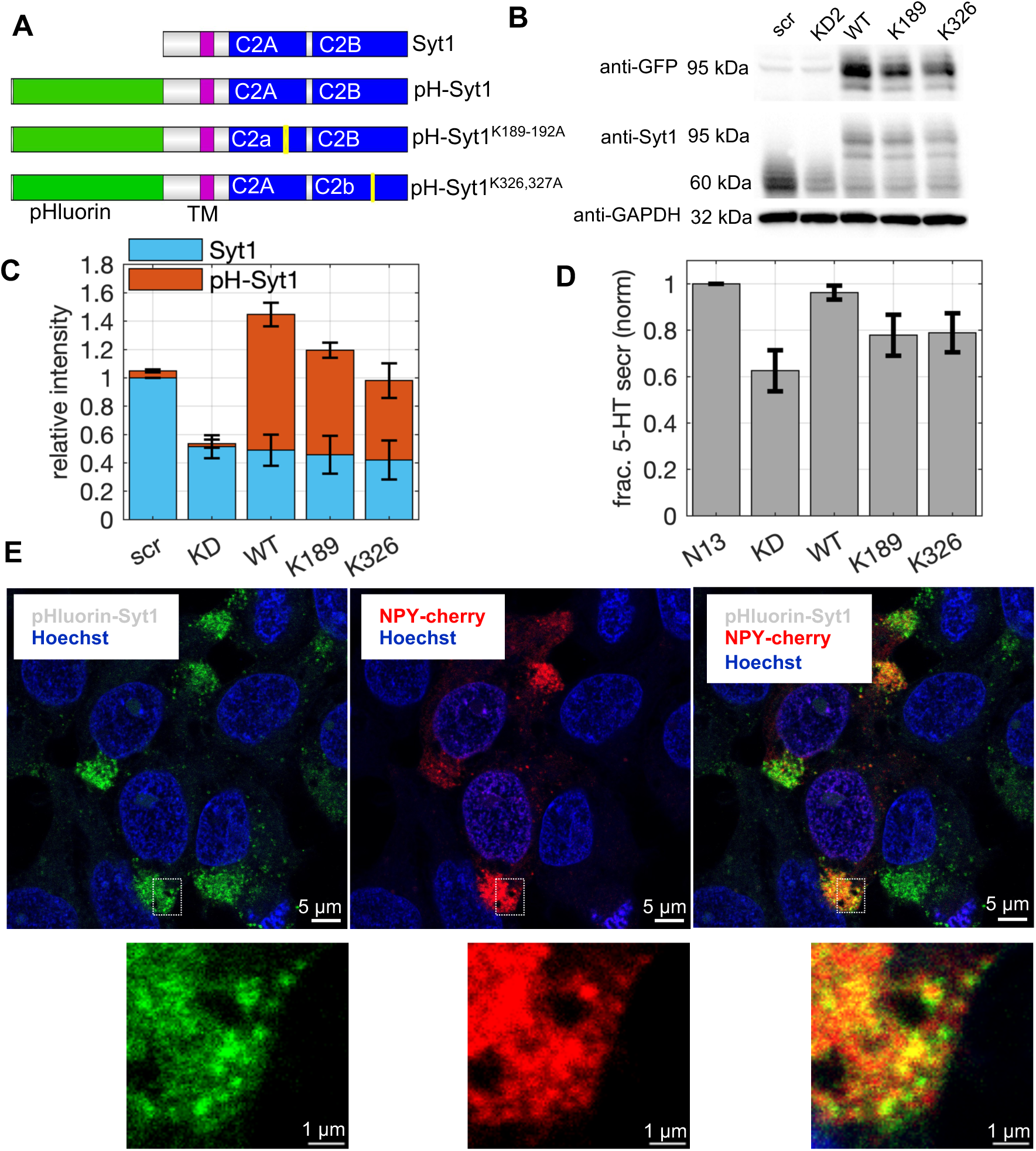
Syt1 is a major sensor for calcium-triggered exocytosis in BON cells. **A**. Domain structures of the Syt1 wild-type and mutant rescue constructs used. **B**. Western blot (WB) analysis of the constitutive expression of *syt1* transgenes in BON cells. A representative result from 3 separate experiments is shown. Scr (scrambled, mock shRNA), KD (knock-down 2), WT (pH-Syt1^WT^), K189 (pH-Syt1^K189–192A^), K326 (pH-Syt1^K326,327A^). The rescue constructs are stably expressed in BON cells constitutively expressing shRNA against *syt1* (KD). The rescue constructs with pHluorin migrate slower, allowing relative amounts of expression from native vs. exogenous loci (see C). **C.** Quantification of the expression of endogenous *syt1* vs *pH-syt1* rescues, from densitometry analysis of blots as in B. (n=3 blots). 50-60 % of endogenously expressed *syt1* is replaced by the expression *of pH-syt1* rescue constructs, for a total amount 0.9-1.45 times the amount in the parental cell line. **D.** Calcium-dependent bulk release of serotonin (5-HT) from BON cells with the genotypes as indicated, from 3 independent experiments. For each genotype, the fraction of total cellular 5-HT released upon stimulation (ionomycin 10 µM, ∼3 s) was calculated, then normalized to release from the parent cell line (src). The error bars represent ± SEM in both C and D. **E.** Exogenously expressed pH-Syt1 is correctly targeted to DCVs. Confocal images of BON cells stably co-expressing the granule marker NPY-mCherry and pH-Syt1 constructs as indicated. After fixation, cells were permeabilized and pHluorin-Syt1 was detected using Alexa488 labeled anti-Syt1 antibodies. The boxed regions are shown at higher magnification below each panel. The Pearson correlation coefficient was coefficient of 0.718 ± 0.037 (SEM, n=4 images), indicating good co-localization (a value of 1 corresponds to perfect colocalization).

To study the roles of Syt1 C2A and C2B polybasic patches, we generated lentiviral expression vectors that stably integrated into the genome *syt1* constructs fused to the pH-sensitive GFP pHluorin^65^. These Syt1 constructs encode the wild-type (WT) sequence, or sequences bearing mutations in the C2A or C2B domains (pH-Syt1^WT^, pH-Syt1^K189–192A^, or pH-Syt1^K326,327A^, Fig. 2A). The rescue constructs fused to pHluorin migrate slower in SDS PAGE gels, allowing the relative amounts of expression from native, knocked down (KD) and rescued expression of WT and point Syt1 mutants to be quantified (Fig. 2B, C). Overall, 50-60% of endogenous Syt1 was replaced with the rescue constructs, for a total Syt1 expression 0.90-1.45 fold the parental BON cell line. Importantly, expression of pH-Syt1^WT^ completely rescued bulk 5-HT release, while pH-Syt1^K189–192A^ or pH-Syt1^K326,327A^ were inefficient in restoring release (Fig. 2D) - consistent with effects on evoked release in cortical neurons, as we reported earlier^28^.

In the acidic milieu of the DCV’s lumen (pH∼5.5), pHluorin fluorescence is largely quenched. Fusion with the PM leads to the rapid neutralization of the luminal pH and a large (6-10-fold) increase of the pHluorin signal, greatly facilitating detection of fusion events^66,67^. In these cells we also stably expressed neuropeptide-Y (NPY), a DCV cargo, fused to mCherry, to facilitate localization of DCVs. While we focus on electrochemical detection of serotonin release here, the dual labeling strategy will be useful in future imaging studies. The rescue constructs were properly localized to DCVs, as pHluorin-Syt1 immunofluorescence and NPY-mCherry signals largely overlapped (Fig. 2E), with a Pearson’s correlation coefficient of 0.718 ± 0.037 (SEM, n=4 images; a coefficient of 1 corresponds to perfect overlap).

In summary, Syt1 is a major calcium sensor for exocytosis in BON cells. Syt7 expression is much lower and could not be detected by immunofluorescence, consistent with a previous report in which Syt7 KD did not affect release from BON cells^61^. The lack of a Syt7-positive DCV pool facilitates interpretation of release studies. Stable expression of pH-Syt1, but not of Syt1^K189–192A^ or pH-Syt1^K326,327A^, rescued bulk release of 5-HT. The inability of the mutants to rescue 5-HT release is not due to a localization defect and must be due to the mutations introduced.

### Both Synaptotagmin-1 C2A and C2B domain polybasic patches are required for efficient DCV exocytosis

Next, we monitored 5-HT release from BON cells at single event resolution using amperometry^68–71^ (Fig. 3). In this approach, catecholamines, serotonin, and other suitable cargo released through exocytosis are oxidized at diffusion controlled rates at the surface of a carbon fibre electrode (CFE) held at +650 mV (vs. Ag | AgCl), above the oxidation potential for 5-HT. The electrode is insulated except at its tip (∼5 µ*m* diameter) that gently touches the cell, minimizing diffusional broadening of the signal. Oxidation transfers two electrons to the CFE from every 5-HT molecule, allowing an oxidation current to be recorded. The rapid oxidation kinetics allow individual release events to be detected as narrow spikes.

**Figure 3.**
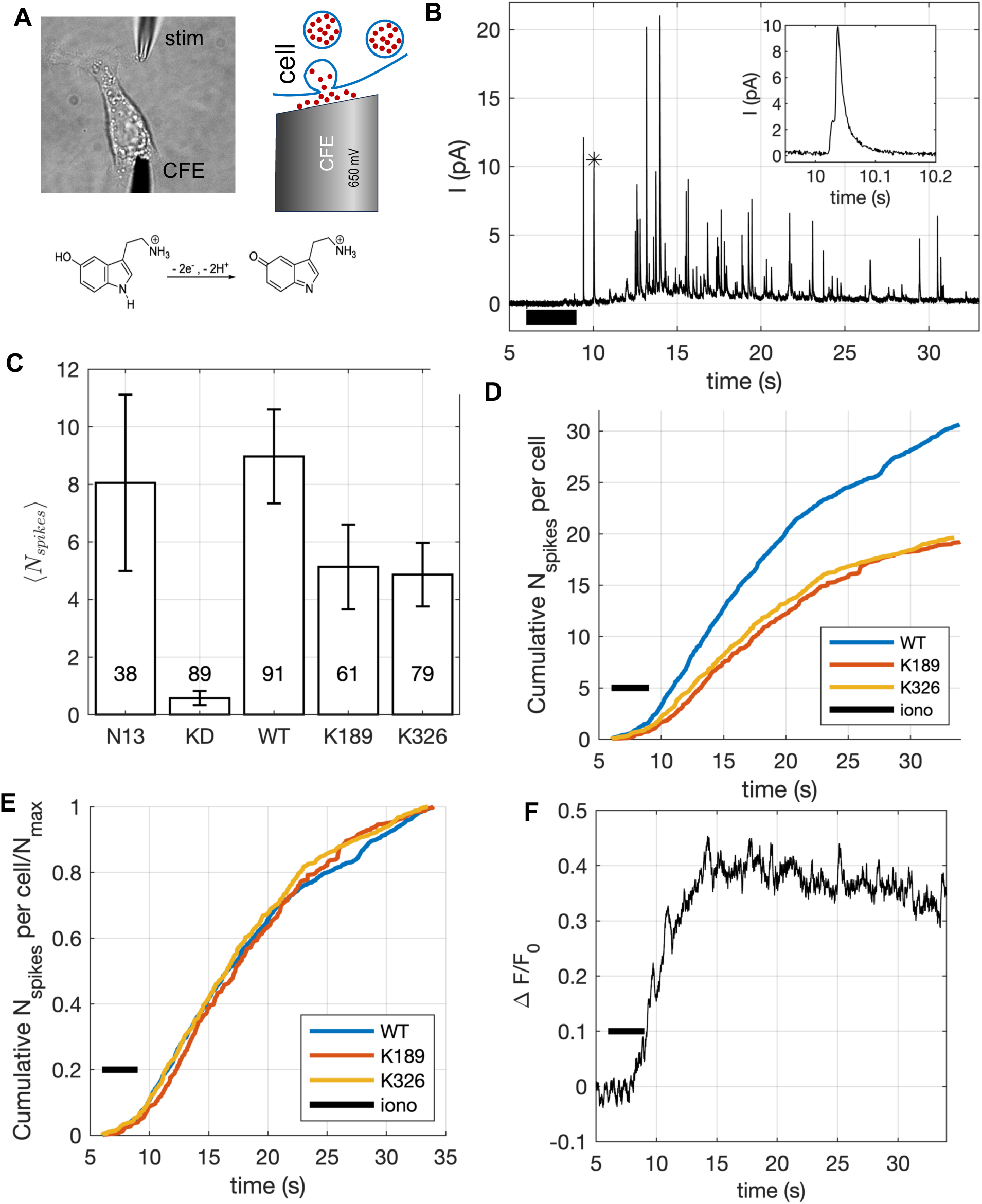
Release of 5-HT is impaired in BON cells expressing pH-Syt1^K189–192A^ or pH-Syt1^K326,327A^. **A**. Depiction of the experiment. A carbon fibre electrode (CFE) held at 650 mV gently touches a BON cell. Stimulation is performed by pressure-driven superfusion of an ionomycin solution from a micropipette placed nearby (“stim”). The principle of detection and the oxidation reaction are shown schematically. **B.** Example of an amperometric trace. The cell was stimulated for 3 s by ionomycin application (black bar). Each exocytosis event results in a brief oxidation spike. The spike marked by an * is expanded in the inset. **C**. Bar plot of the total number of spikes (to t=23 s) averaged over the number of tested cells, including cells that did not respond to stimulation (<5 spikes). N13 is the parent BON cell line, the other symbols are as in Fig. 2. Error bars represent S.E.M. The number of cells tested are indicated above every bar. **D**. Release kinetics from experiments as in B, plotted as the cumulative number of spikes per cell. Only cells that responded to stimulation (≥5 spikes) were included. Syt1 KD impairs release, pH-Syt1-NPYmCHerry restores release. Rescue with pH-Syt1^K189–192A^ or pH-Syt1^K326,327A^ are both defective, especially in light of remaining endogenous WT Syt1. **E.** Release kinetics as in D, normalized to the maximum number of spikes per cell for every group. There is no detectable delay in release. **F**. An example of changes induced in intracellular calcium upon stimulation using the calcium indicator Fluo-4. The parental, unlabeled BON N13 cells were used for these experiments to avoid overlap with other fluorescent molecules expressed in the rescued cell lines, but otherwise the conditions were the same as for D.

BON cells normally store low amounts of 5-HT. To improve signals, we incubated BON cells in media containing 5-HT^59,60,64^. Following previous work^59,60,64^, to stimulate release, we used ionomycin, a calcium ionophore that elevates intracellular calcium, bypassing calcium-channels, as BON cells respond poorly to stimulation by depolarization^54^. We placed a CFE gently on a BON cell, and after first recording a baseline for 6 s, stimulated release by pressure-driven superfusion of a solution of ionomycin for 3 s through a micropipette placed ∼10 µ*m* away (Fig. 3A). For BON Syt1 KD cells rescued with pH-Syt1^WT^, stimulation caused many oxidation peaks to appear for ∼30 s, as shown for an example in Fig. 3B.

To compare the total amount of 5-HT released, we calculated the average number of cumulative spikes per cell up to 23 s after the start of the recording (Fig 3C). In this analysis, we included all cells tested, including those that did not respond to stimulation. Syt1 KD reduced release ⪎10-fold, while rescue with pH-Syt1^WT^ fully rescued it. Syt1^K189–192A^ or pH-Syt1^K326,327A^ rescued release partially, consistent with bulk 5-HT release measurements (Fig 2D).

To compare release kinetics among cells expressing pH-Syt1^WT^, Syt1^K189–192A^, or pH-Syt1^K326,327A^, we plotted the cumulative number of spikes for all cells, divided by the number of cells that responded to stimulation (≥ 5 spikes within 34 s of recording) in Fig. 3D. Consistent with the results above, release from cells expressing Syt1^K189–192A^ or pH-Syt1^K326,327A^ was impaired compared to cells expressing pH-Syt1^WT^. When the cumulative number of spikes was normalized to the maximum for every condition, we could not detect any differences in release kinetics (Fig. 3E).

We monitored changes in intracellular calcium induced under identical conditions using the parental, unlabeled BON cells, loaded with Fluo-4, a fluorescent calcium indicator^72^. We found that [Ca^2+^]_i_ rose within 2-5 s to its half maximum after the end of the ionomycin application (Fig. 3F). Thus, the kinetics of release may be limited by the kinetics of elevation of intracellular calcium by the ionomycin application.

To surmise the results so far, the efficient calcium-stimulated release of serotonin from BON cells requires both C2 domains of Syt1. We could not detect any difference in release kinetics among cells expressing Syt1^WT^, Syt1^K189–192A^ or pH-Syt1^K326,327A^, but this may be due to the slow rise of intracellular calcium with ionomycin application.

### Synaptotagmin-1 C2A and C2B domain polybasic patches contribute to DCV docking

We next examined DCV docking in BON cells expressing pH-Syt1^WT^, pH-Syt1^K189–192A^, or pH-Syt1^K326,327A^ using electron microscopy (Fig 4A-C). Cells were fixed under resting conditions or after a ∼3 s stimulation by ionomycin, mimicking the conditions used for amperometry. They were stained, resin embedded, sectioned, post-stained, and imaged by transmission electron microscopy (EM). DCVs near the plasma membrane were identified through their characteristic dark, electron-dense core. Consistent with previous reports, some DCVs were lighter and some were non-spherical^53,61,62^. To compare DCV sizes, we traced outlines of the DCVs, and calculated the area. We found no significant differences among the groups (Fig 4D), suggesting that the different Syt1 mutants do not affect DCV size. We then calculated the nearest distance between a DCV and the plasma membrane, *d*. In resting cells expressing pH-Syt1^WT^, there was a higher density of DCVs with *d* < 25 nm compared to those expressing pH-Syt1^K189–192A^, or pH-Syt1^K326,327A^ (Fig 4E). Upon a 3s stimulation, a portion of this docked population was lost in cells expressing pH-Syt1^WT^, likely through exocytosis. For cells expressing Syt1 mutants, the distribution of *d* did not change significantly with stimulation (Fig 4F). The simplest interpretation is that upon stimulation, DCVs had to mobilize from *d* > 25 nm to undergo fusion with the plasma membrane in these cells.

**Figure 4.**
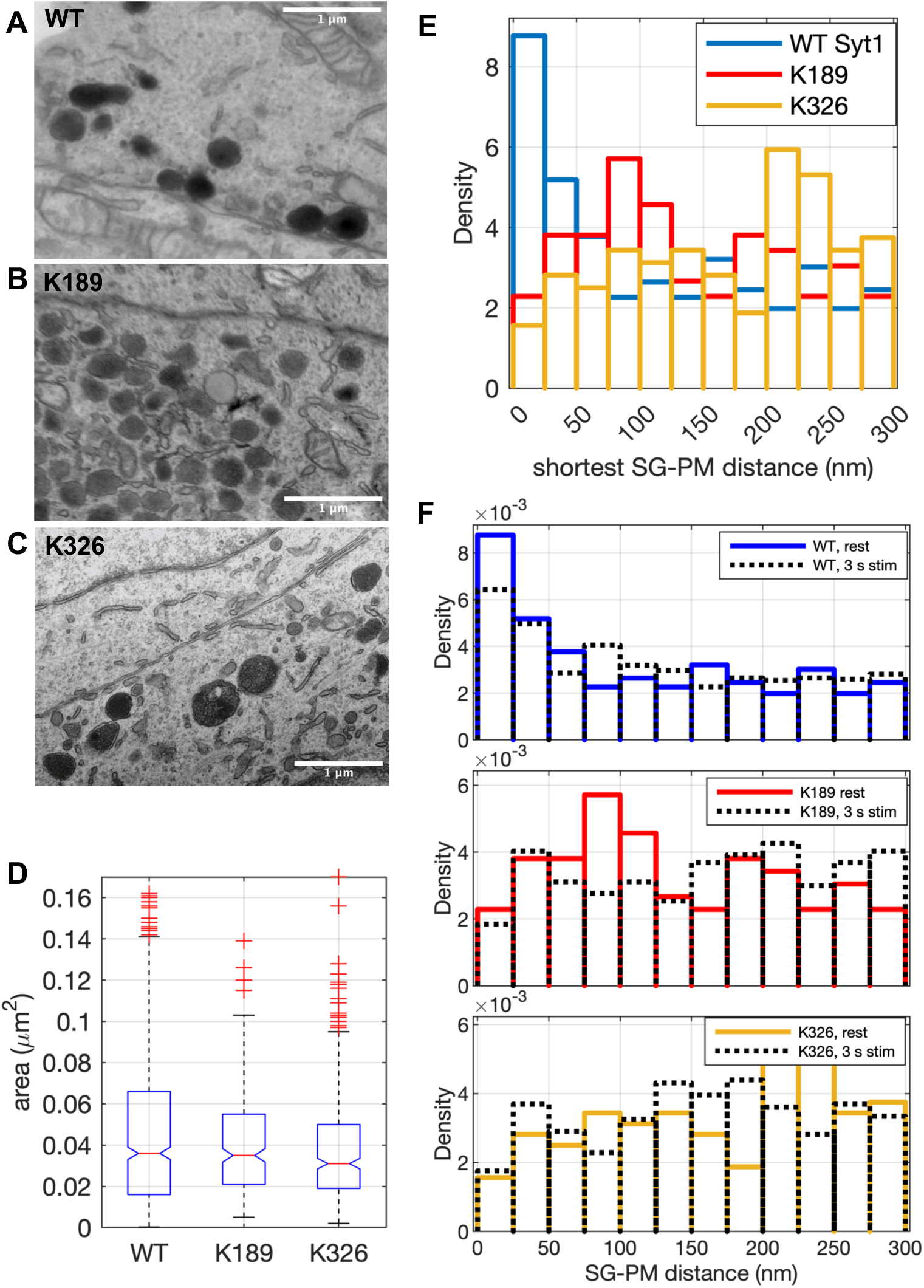
Syt1 polybasic patches contribute to DCV docking to the plasma membrane. **A**. An electron micrograph of Syt1 KD BON cells rescued with pH-Syt1^WT^ (WT). **B-C**. Same, for rescue with pH-Syt1^K189–192A^ (K189, B) or pH-Syt1^K326,327A^ (K326, C). **D.** Comparison of DCV areas among the groups (*p* = 0.57 for the null hypothesis that the data in each group comes from the same distribution, using the Kruskal-Wallis test). **E.** Distributions of shortest DCV-PM distances *d*, for *d* ≤ 300 nm, for non-stimulated cells, for the groups shown. Labels are the same as in A-C. **F**. Changes in the DCV-PM distances upon a brief, ∼3s stimulation. The distributions before stimulation are the same as in E, replotted for easier comparison. Samples were prepared from at least two independent cultures.

In summary, Syt1 is important for DCV docking in BON cells, consistent with its previously reported role in DCV docking in mouse adrenal chromaffin cells^20,23^. The polybasic patch in both C2 domains play a role in docking, suggesting both domains interact with the PM. The role of C2B is consistent with previous reports in neurons^16,17^, but the role of C2A is newly identified, to the best of our knowledge.

### Synaptotagmin-1 C2A and C2B domain polybasic patches counter fusion pore expansion

Amperometry allows single fusion events to be examined with high sensitivity and temporal resolution, providing information about fusion pore properties. The parameters that can be quantified from an isolated amperometric event are shown schematically in Fig. 5A. For a subset of spikes, an initial shoulder called the pre-spike foot (PSF) signal can be detected, with the associated duration *t_PSF_* and amplitude *I_PSF_* The PSF reflects flux of oxidizable cargo through the initial fusion pore whose size is ≲ 1 nm^44,46,47^. The signal then rises rapidly to reach a maximum amplitude *I_max_* before decaying more slowly with a width at half-amplitude *t_1/2_* The spike’s rise and decay reflect the flux through the fusion pore of the cargo that is detected at the electrode surface. The flux is affected by the fusion pore size and the cargo concentration remaining in the fused DCV. Though larger to support the additional flux, this pore may still be only a few nanometers in diameter^73–78^. The integral of the amperometric current is the charge *Q*_0_, which is related to the number of molecules oxidized at the electrode surface via Faraday’s law, *Q*_0_ = *nFN*, where *N* is the number of molecules oxidized, *F* is the Faraday constant, and *n* is the number of electrons exchanged at the electrode^69^ (*n* = 2 for 5-HT).

**Figure 5.**
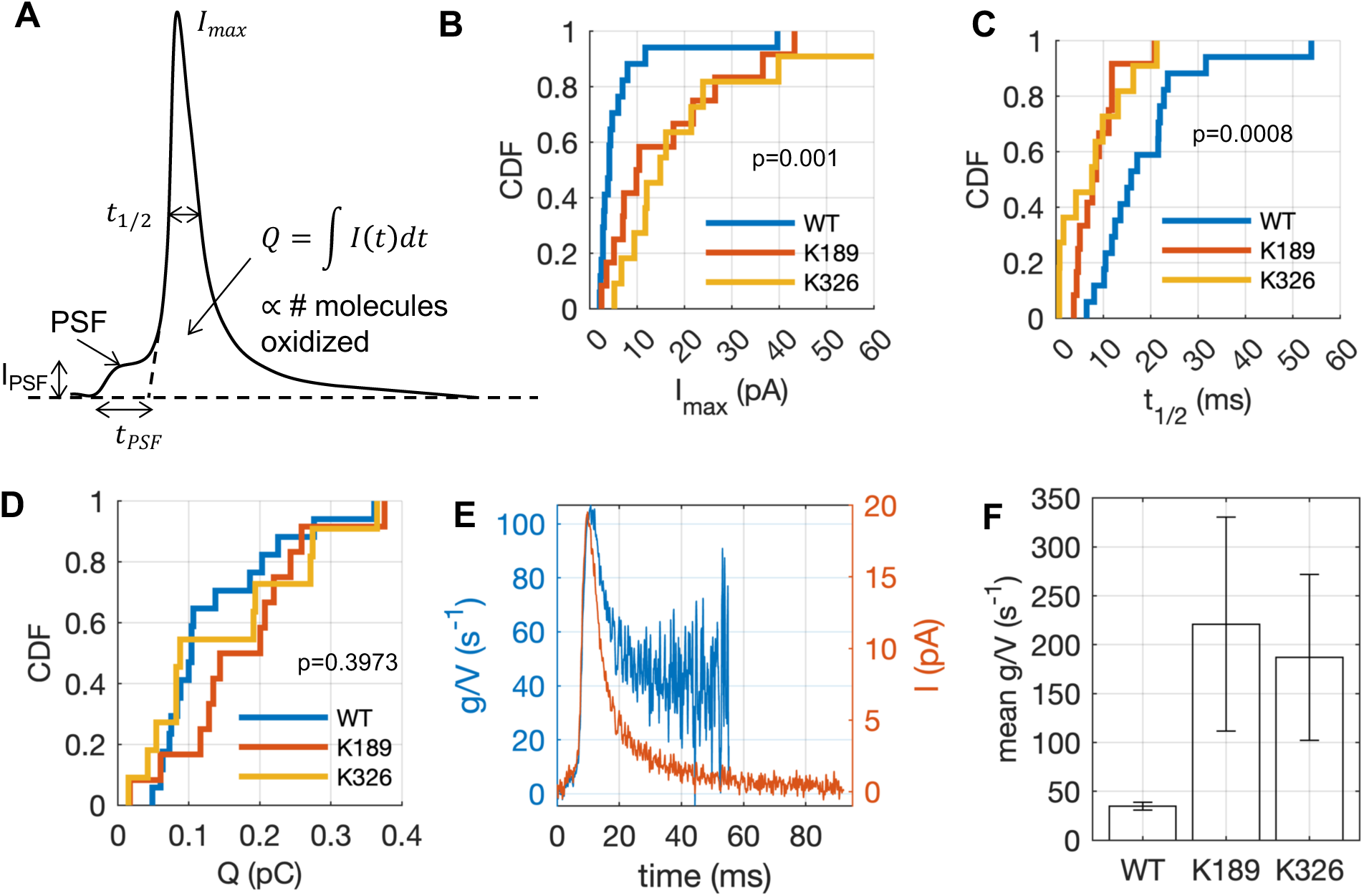
Syt1 polybasic patches control fusion pore permeability. **A**. Schematic of a single amperometric oxidation event (see Fig. 3B). **B**. Cumulative distribution function (CDF) of maximum spike amplitudes averaged over BON cells expressing Syt1^WT^(WT), Syt1^K189–192A^ (K189), or Syt1^K326,327A^ (K326). Syt1^WT^ spikes have lower amplitude on average. See Table 1 for a summary. **C**. Cumulative distribution of spike widths at half amplitude for BON cells expressing Syt1^WT^, Syt1^K189–192A^, or Syt1^K326,327A^ (symbols as in B). Syt1^WT^ spikes last longer. **D.** Cumulative distribution of oxidation charges for individual events for BON cells expressing Syt1^WT^, Syt1^K189–192A^, or Syt1^K326,327A^. No significant difference is found, implying the same amount of 5-HT is released per event for the different conditions. The *p*-values in B-D are returned from the Kruskal-Wallis test for the null hypothesis that all data come from the same distribution. **E.** Example of a trace of fusion pore permeability scaled by DCV volume (*g*/*V*) as a function of time (blue trace). The corresponding amperometric current *I*(*t*) is shown in red. The integral is carried only to 60% of the total spike duration (see text and Methods). **F**. Mean *g*/*V* for 0-60% of spike duration, averaged over cells in each group. Error bars represent SEM. For WT, K189, and K326 groups, the number of cells (spikes) were 17 (197), 12 (108), 11 (131), respectively. Mean values for every cell were averaged across cells. *p* = 0.003 and 0.020 for the mean ranks of K189 and K326 data compared against WT (see methods).

**Table 1.** Amperometric spike parameters. Mean values for every cell were computed first, then the cell means were averaged over the number of cells^180^. The errors are ± SEM.

| | $\langle I_{max} \rangle$ (pA) | $\langle t_{1/2} \rangle$ (ms) | $\langle Q \rangle$ (C) | $N_{\text{molecules}}$ | $N_c(N_s)$ |
| --- | --- | --- | --- | --- | --- |
| pH-Syt1<sup>WT</sup> | $6.58 \pm 2.34$ | $18.34 \pm 2.91$ | $0.1517 \pm 0.0383$ | $(7.86 \pm 1.99) \times 10^5$ | 17 (197) |
| pH-Syt1<sup>K189-192A</sup> | $20.82 \pm 7.54$ | $8.27 \pm 1.78$ | $0.2264 \pm 0.0694$ | $(11.7 \pm 3.60) \times 10^5$ | 12 (108) |
| pH-Syt1<sup>K1326,327A</sup> | $15.99 \pm 7.01$ | $10.27 \pm 2.64$ | $0.1748 \pm 0.0507$ | $(9.06 \pm 2.63) \times 10^5$ | 11 (131) |

Interestingly, spikes recorded from BON cells expressing pH-Syt1^WT^ had lower amplitude on average, ⟨*I_max_*_’_⟩ = 6.58 ± 2.34 pA), compared to those from cells expressing pH-Syt1^K189–192A^ (20.82 ± 7.54 pA) or pH-Syt1^K326,327A^ (15.99 ± 7.01 pA), as shown in Fig. 4B and Table 1. Cells expressing pH-Syt1^WT^ also had longer spike durations (*t_1/2_* = 18.34 ± 2.91, 8.27 ± 1.78, and 10.27 ± 2.64 pA, respectively, for cells expressing pH-Syt1^WT^, pH-Syt1^K189–192A^, and pH-Syt1^K326,327A^ (Fig. 5C and Table 1). By contrast, there was no significant difference in the charge, *Q*_0_, among the three groups of cells, indicating that on average, the same amount of 5-HT was released per fusion event, corresponding to about a million molecules and is in good agreement with a previous report^59^ (Fig. 5D and Table 1). For PSF parameters, no significant differences were observed.

Overall, these results show that Syt1 polybasic patch mutations cause faster cargo release during individual fusion events, implying a larger pore permeability. To test if this were indeed the case, we calculated the pore permeability *g*, scaled by vesicle volume *V*, for non-overlapping amperometric events, following Jackson^75^:

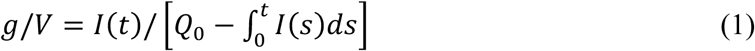

Here the denominator is proportional to the number of molecules remaining in the vesicle at time *t*, with *Q*_0_ the total charge (integral of *I*(*t*) from start to end of an amperometric event). The only assumption in deriving eq. (1) is that the flux through the fusion pore is proportional to the cargo concentration within the vesicle, valid if the flux through the fusion pore is rate limiting (implying negligible concentration gradients inside the vesicle and a low concentration outside)^75^.

An example of a (volume-scaled) fusion pore permeability as a function of time for a single event is shown in Fig. 5E (blue trace). The amperometric current profile, *I*(*t*) is also shown (red trace, right axis). Because the denominator in eq. (1) tends to zero, *g*/*V* becomes increasingly noisy toward the end of the event. For this reason, only the initial 60% of current traces were converted to *g*/*V*. Interestingly, as was observed by Jackson for release from mouse chromaffin cells, pore permeability reached a plateau after going through a peak that was slightly delayed compared to the peak in *I*(*t*). That is, after a transient, the pore reached a size that remained stable (or fluctuated rapidly around a stable size) for at least tens of milliseconds. The subsequent fate of the pore cannot be known with amperometry, because most of the cargo has left the vesicle by this time and little or no amperometric signal remains^46,75^. However, imaging studies using total internal reflection fluorescence (TIRF)^63,79^, confocal^80,81^, or superresolution microscopy^82^ are consistent with long-lived fusion pores, though pore size is harder to estimate, and dynamics cannot be probed with high time resolution. For most events that we recorded from BON cells, *g*/*V* followed a typical time-course as shown for the example in Fig. 5E, but some events were characterized by a slow rise to a stable plateau without going through a peak. The mean *g*/*V* for the initial 60% of an amperometric event, averaged over all cells for the different experimental groups is plotted in Fig. 5F. The quantity *g*/*V* was 4-5 fold larger for cells expressing Syt1 mutants than for WT. This effect was not due to a smaller vesicle volume for the mutants, as we found no difference in DCV size by EM analysis (Fig. 4D). Thus, the polybasic patches in both C2 domains contribute to maintaining a small fusion pore.

## DISCUSSION

### DCV docking by Syt1 C2A and C2B polybasic patches

Syt1 is a major calcium sensor for SV and DCV exocytosis^1^. Most past work focused on understanding the role of Syt1 in rapid calcium-triggering of SV or DCV release. However, Syt1 also plays important roles in vesicle docking and fusion pore regulation, which have received relatively less attention. Perturbations to Syt1 function can affect the release process at the docking, fusion, or post-fusion stages, or a combination of these. Thus, it is important to understand how these functions relate to one another, and which stage(s) of exocytosis are affected when Syt1 function is perturbed.

For SV and DCV exocytosis, vesicles are first delivered to the vicinity of the PM where they undergo “priming” to acquire fusion-readiness, Fig. 6. Morphologically, priming involves at least two sequential states, “tethered” and “docked” in which the vesicle-PM distance, *d*, is different^3,32,33,35,36^. Biochemically, it requires activation of large priming factors such as Munc13 or CAPS by Ca^2+^, diacylglycerol (DAG), or other components^3,32,33,35,36,83^. Munc13 is ∼30 nm long, and can tether vesicles within this distance^3,32,36^. Munc13 and/or CAPS, together with Munc18 catalyze the formation of SNARE complexes between the vesicular v-SNARE VAMP2/Synaptobrevin-2, and the PM t-SNAREs Syntaxin-1 (Stx1) and SNAP25, or related isoforms^2,3,84–86^. Formation of SNARE complexes drives membrane fusion, but under resting [Ca^2+^]_i_ (∼100 nM), fusion is inhibited by Syt1 (and possibly complexins) by mechanisms that are not completely resolved^2–4,87–90^. It is thought that in the primed state SNARE complexes are at least partially assembled^86^, but the degree of SNARE domain assembly is debated, from minimally assembled SNAREs (with only membrane distal N-termini interacting, ∼8 nm) to fully zippered ones (≲5 nm)^36^. In addition to SNAREs, Syt1 is also implicated in DCV^20^ and SV^16,17^ docking, with *d* ≲12 nm^17^. Thus, priming involves a shortening of the vesicle-PM distance *d*, from 20-30 nm in the tethered state to 3-8 nm in the primed state^36^.

**Figure 6.**
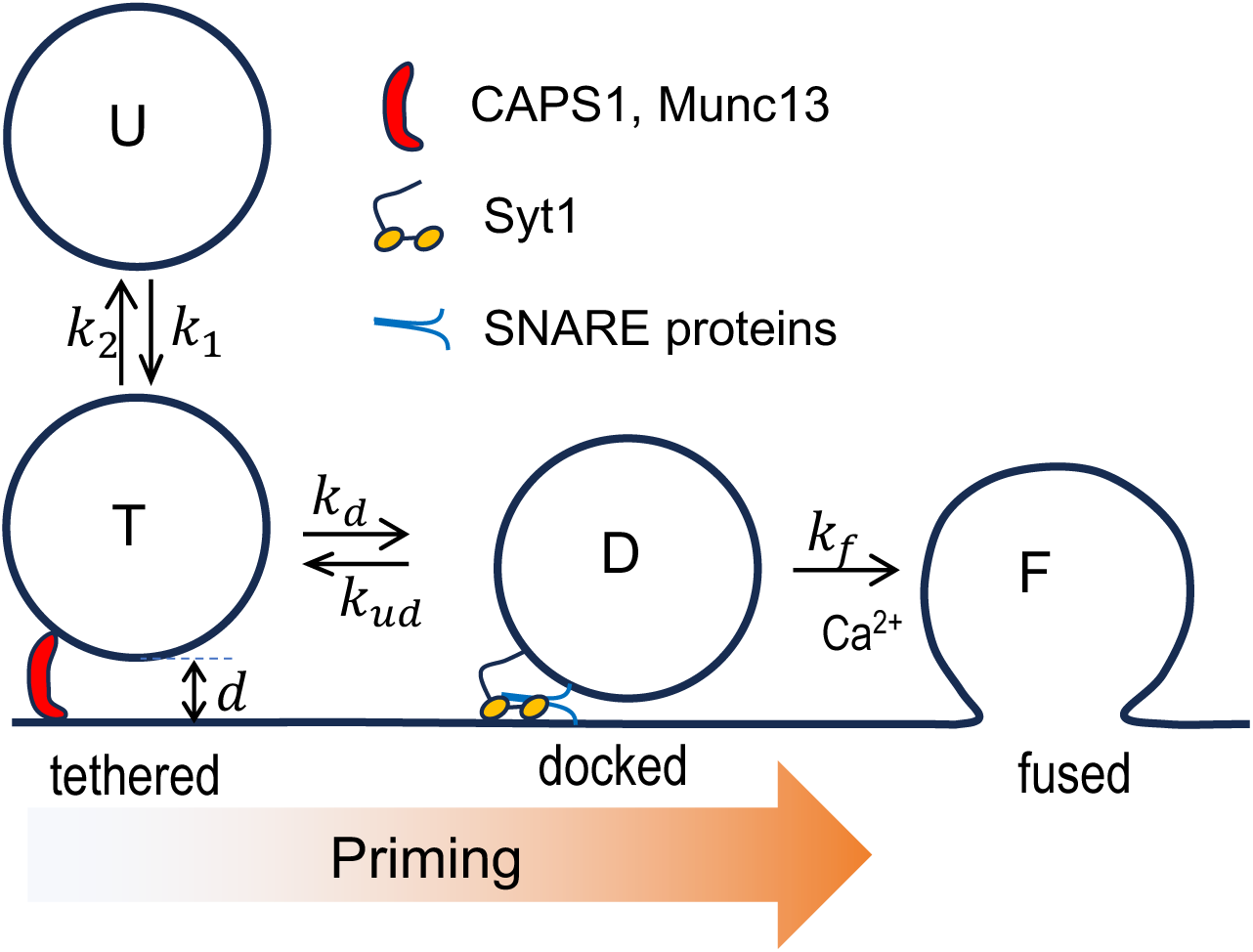
Model of two-stage priming in BON cells. Undocked (U) DCVs reversibly tether (T) to the PM. Initial tethering is mediated by large tethering molecules such as CAPS and/or Munc13, which also assist subsequent stages of DCV maturation at the PM. Docked DCVs (D) are closer to the PM. Docking involves Syt1 and likely SNARE proteins and acidic lipids in the inner leaflet of the PM. An elevation of [Ca^2+^]_i_ leads to fusion (F). Release kinetics and extent depend on the amount of docked DCVs. If the docked population is sparse, the release rate may be limited by the rate of docking.

Although the model depicted in Fig. 6 has been considered for DCV priming for some time^62^, a similar model has only recently been applied to SV priming^32,34–36^. This minimal two-stage priming model explains aspects of synaptic plasticity^32,34–36^. It also suggests the roles previously assigned to the fusion step may in fact be upstream^91,92^. However, testing the model is particularly difficult for SV exocytosis, because monitoring SV docking is challenging, and many neurons express multiple Syt isoforms^93–95^. By contrast, DCV docking defects monitored using electron microscopy lead to much clearer phenotypes^19,20,22,96^ than for SVs^16,17,97^, due to the sparser packing and lack of tethering among DCVs. DCV docking can also be studied using TIRF microscopy; DCV-PM attachment is evident as a sudden drop in DCV mobility both in the imaging (*xy*) plane and the plane perpendicular to it (*z*) in BON^19,22,49,54,62^ and chromaffin^50^ cells (but see^98^). In some cases, distinct DCV-PM attachment states could be detected^62,99^. We found that about half the DCVs in BON cells made a ∼20 nm step toward the PM after stimulation^62^, corresponding to the ***T*** → ***D*** transition in Fig. 6, but the underlying molecules were not identified. Our current results implicate Syt1 as a strong candidate in this transition.

Syt1 couples Ca^2+^ influx to membrane fusion via its two C2 domains that bind Ca^2+^, acidic lipids, SNAREs, and other effectors^100–102^ (Fig. 1A). How Ca^2+^ binding to Syt1 leads to fusion is debated^103–111^. Syt1 KO produces docking defects in neurons^112,113^ and neuroendocrine cells^114^, indicating Syt1 is needed for docking of DCVs and SVs. However, since synchronous release is lost in Syt1 KO neurons, relating release to the vesicle-PM distances *d* was not possible. The Syt1 C2B polybasic patch is required for tight docking of SVs^108,113^. Partial neutralization of the patch (K325, 327A) produced docking defects that correlated with reduced evoked release^108^.

Previously, we found that Syt1^K326,327A^ in cortical neurons led to impaired evoked inhibitory post synaptic currents (eIPSCs) with a slight delay in reaching the eIPSC peak^115^, with no significant effect in spontaneous release or the RRP. Thus, the K326,327A mutation reduces the probability of release P_*r*_during a brief action potential. Neutralization of Syt1 C2A polybasic patch (Syt1^K189–192A^) led to similar, but slightly less severe release defects^115^. Importantly, a docking defect may not be evident in RRP measurements^91,92^, leading to the incorrect conclusion that P_*r*_ from the D state is affected, while the true effect may be a smaller ***D*** pool. Indeed, here we found that Syt1^K189–192A^ and Syt1^K326,327A^ led to a smaller docked DCV pool in BON cells under resting conditions, which correlated with those in serotonin release evoked by a short stimulation.

The docking defects we identified here for Syt1 mutations are consistent with the kinetic delay in cortical neurons we observed previously^28^. A kinetic delay may also be present in BON cells expressing the Syt1 mutants, but may be masked by the slow elevation of [Ca^2+^]_i_ we elicit using ionomycin stimulation. We plan to explore release kinetics with better time resolution in future studies, using UV-uncaging of calcium.

How does Syt1 mediate vesicle docking? A likely PM binding partner is the Stx1-SNAP25 t-SNARE acceptor complex^114^, possibly with assistance from Snapin^116–122^. Structures of Syt1-SNARE complexes are consistent with Syt1-t-SNARE interactions^123,124^. Yet, other work argued that Syt1 does not bind to SNARE complexes under physiological conditions^125^. Two groups concluded that Syt1 docks vesicles by directly binding PI(4,5)P_2_ on the PM^113,126^, while another concluded this is not the case^127^. Because there are several copies of Syt1 on a DCV or SV, it is possible that multiple weak interactions between Syt1 and the t-SNAREs add for stably docking vesicles to the PM *in vivo*. Alternatively, Syt1 may use coincidence detection, simultaneously binding PI(4,5)P_2_ and t-SNAREs. Interestingly, our results suggest that the Syt1 C2A domain interacts with the PM, not the vesicle membrane, at least during the docking stage.

### Modulation of fusion pore dynamics by Syt1 polybasic patches

Upon elevation of [Ca^2+^]_i_, primed vesicles fuse with the PM^2,84,86,128^. This last step minimally requires the neuronal SNARE complex, Syt1, and Cpx1^4,6,89^. The initial merger of the membranes results in a narrow fusion pore ∼1-3 nm in diameter^44,46,47,129^. Pore dynamics determine release kinetics and the mode of vesicle recycling. The pore can fluctuate in size, flicker open-closed and reseal after partial release of cargo or dilate fully^44,46,47,129^. Because DCVs contain a range of cargoes, the pore can act as a molecular sieve, controlling what is released^130–134^. Fusion pore dynamics also affect release of neurotransmitters and SV _recycling_135–142_. SNARE proteins_37,76,77,129,143–148_, Synaptotagmins_37–39,41–43,149–153_, and_ lipids^134,154–156^ affect pore dynamics but the mechanisms are poorly understood. Notwithstanding, a number of key factors that modulate SNARE number or proximity to the PM have been identified. Increased number of SNARE proteins at the fusion site increase pore permeability^157–161^ by crowding at the pore’s waist^161^. Flexible linkers inserted between the SNARE and the TMDs in VAMP2, or truncation of SNAP25, slow pore expansion^162–164^. Syts also affect fusion pores^103,165–173^ but mechanisms are even less clear. In chromaffin cells where Syt1 and Syt7 are sorted to distinct DCV pools, fusion pores of Syt7 DCVs dilate more slowly^172^.

Here, we found that neutralization of either the C2A or C2B domain of Syt1 resulted in larger fusion pores during fusion of DCVs. These larger pores released the same amounts of cargo, but faster, during individual fusion events. There are at least three plausible mechanisms. First, Syt1 can cooperate with the SNARE complex to exert a calcium-dependent mechanical lever action to expand the distance between the fused membranes^37^. This incurs a bending energy penalty that is offset by expanding the pore^37^. The lever action is driven by the rotation of the Syt1 C2B-SNARE complex upon calcium binding to the C2B domain which induces its reorientation with respect to the PM. However, this model does not account for the effects of the C2A domain.

Second, the fusion machinery might be less tightly organized with the Syt1 mutants, leading to a faster expansion of the fusion pore. Indeed, Syt1 has been proposed to form a washer-like ring between an SV and the PM that dissociates upon elevation of [Ca^2+^]_i_^6^. Finally, Syt1 may affect fusion pore dynamics *via* lipid signaling. In stimulated cultured hippocampal neurons, Syt1 interacts with the phosphatidylinositol 4-phosphate 5-kinase type Iγ (PIP5K1C) which locally produces PI(4,5)P_2_ from phosphatidylinositol 4-phosphate (PI(4)P), facilitating endocytosis^174^. Interestingly, local PI(4,5)P_2_ production was also observed in insulin-secreting cells at sites of exocytosis^134^. Increased local PI(4,5)P_2_ production required PIP5K and facilitated recruitment of endocytic factors, including BAR domain proteins amphiphysin, SNX9, and endophilin, which in turn constricted fusion pore expansion^134^. Although it is not yet clear if Syt1 interacts with PIP5K in DCV exocytosis, and when the interaction occurs in neurons, it is possible that Syt1’s roles in endocytosis and on fusion pore regulation are coupled *via* lipid signaling. If this is the case, our results would then imply that the neutralization of the polybasic patches in Syt1 C2A and C2B regions disrupt this signaling. This in turn would impair the recruitment of endocytic factors to the fusion pore’s neck that stabilize it, and thereby lead to faster dilation of the fusion pore.

In summary, we found that neutralization of the polybasic patches in either of Syt’s1 two C2 domains leads to correlated docking and release defects. The same mutations also lead to larger fusion pores, suggesting that the docking, fusion triggering, and post-fusion roles of Syt1 are not independent.

## MATERIALS AND METHODS

### Cell lines

The original BON cell line was not clonal^53^. We used a clone named N13, generated and kindly provided by Bruno Gasnier (CNRS, Université Paris Cité, France)^175^. BON N13 cells were cultured in DMEM/F12 medium containing 10% heat in-activated FBS and 1X of Pen-Strap antibiotics (15140122, Thermo Fischer Scientific, Waltham, MA), 1X of non-essential amino acids (11140050, Thermo Fischer Scientific) and 1 mM of sodium pyruvate. Cells were grown in a cell incubator at 37°C under 5% CO_2_.

#### Lentiviral shRNA and generation of pL309-pH-rSyt1 PGK-NPY-mCherry

Human Syt1 gene shRNA pLKO lentiviral plasmids TRCN0000053856 and TRCN0000053857 were obtained from the Yale Cancer Center Genome-wide Mission human shRNA lentiviral library collection (Millipore Sigma). The shRNA target sequences on these plasmids are: GCGACTGTTCTGCCAAGCAAT and GCAAAGTCTTTGTGGGCTACA respectively. The pL309 lentiviral plasmids used to express pH-rSyt1 and NPY-mCherry was a gift from T. Südhof^176^, and the rat Syt1 cDNA tagged with GFP-phluorin (pcDNA3-pH-rSyt1) was generated by the V. Haucke laboratory^65^. First, we cloned the sspH-TEV-rSyt1 fragment from pcDNA3-pH-rSyt1 into pL309 as a BamHI-EcoRI fragment to generate pL309-pH-rSyt1. Then, the human PGK promoter with the NPY-mCherry was cloned into the EcoRI-BsrGI sites to generate pL309-pH-rSyt1 PGK-NPY-mCherry plasmid. The Syt1 C2A and C2B mutated sequence fragments were ordered as gBLOCK fragments from IDT and cloned into pL309-pH-rSyt1 PGK-NPY-mCherry that were cut with AgeI-EcoRI to replace the WT Syt1 DNA with the mutated fragments. The mutated fragments were sequenced to verify that they contained the mutations.

Production of the lenti-viral particles and infection of BON-N13 cells was done as described before^177^.

### Western blotting

To analyze Syt1 depletion efficiency and expression of pH-rSyt1 constructs, stably infected BON N13 cells with the different constructs were trypsinized, counted and 2 mL of 100,000 cells/mL suspension for each cell line was plated into 6-well plates and incubated 48hr at 37°C under 5% CO_2_. Total cell lysates were prepared by moving the 6-well plates into an ice bucket, removing the media, washing the wells with 1mL of ice-cold 1X PBS and adding 350 µL of ice-cold RIPA (Thermo Fisher Scientific, Waltham, MA) buffer with 1X protease inhibitor (EDTA-free, Roche) and 1mM PMSF. The cells were then scraped and resuspended from the wells with a 1000 µL pipette tip and transferred to a 1.5 mL ice-cold microcentrifuge tube. Lysates were centrifuged at 14,000 xg for 10 min at 4°C. The supernatant was then transferred to a new ice-cold 1.5mL microcentrifuge tube and store at −20°C. A 25 µL of total lysate mixed with 6 µL of 4X loading buffer per sample was incubated at 37°C for 10 min before loading the samples into a 10% SDS-PAGE gel to separate the proteins at 140 V for 1.5 hr. The gel was then transferred into a nitrocellulose membrane at 120 V for 1 hr at 4°C.

The membrane was blocked in 5% (w/v) milk in PBS-T (PBS with 0.1% (v/v) Tween-20) for 60 min. Molecular weight regions above 50 kDa were cut from the membrane and incubated overnight at 4°C with primary antibodies against Syt1 (cat# 105 011, Synaptic Systems, Goettingen, Germany) and the region below 50 kDa was incubated with anti-GAPDH (cat#2198, Cell Signaling, Danvers, MA) in blocking solution. Next day, the membranes were washed three times for 10 min each with PBS-T and then incubated with HRP-conjugated secondary antibody in blocking solution for one hour, washed again three times with PBS-T for 10 min each and incubated for 5min with SuperSignal West Pico PLUS Chemiluminescent Substrate (Thermo Fisher Scientific) before detection. The membrane incubated with Syt1 antibody was washed with PBS-T twice and incubated for 15 min with Restore Western Blot Stripping Buffer (Thermo Fisher Scientific). After the stripping incubation the membrane was washed twice with PBS-T and incubated for 1hr in blocking solution. The membrane was re-blotted with anti-GFP antibody (Thermo Fisher Scientific, cat# A-11122) over-night at 4°C in blocking solution.

### Immunofluorescence

Transfection of HEK293T cells with plasmid pCMV-ratSyt7 (kindly provided by J.E. Rothman) was carried out using Lipofectamine 3000 (L3000001, Thermo Fisher Scientific) according to the manufacturer’s instructions. BON or HEK293T cells were cultured on poly-L-lysine-coated coverslips, fixed with 4% paraformaldehyde (PFA) for 10 min at room temperature, and permeabilized with 0.1% saponin in PBS for 20 min. Following permeabilization, the cells were blocked with 1% (w/v) bovine serum albumin in PBS for 30 min. They were then incubated overnight at 4°C with either mouse anti-Syt1 antibody (1:1000; 105.011, Synaptic Systems) or mouse anti-Syt7 antibody (1:1000; MA5-27654, Synaptic Systems,). The following day, cells were washed and incubated for 1 hour at room temperature with Alexa Fluor 488-conjugated anti-mouse antibody (1:1000; A28175, Thermo Fisher Scientific). Nuclei were stained with Hoechst 33342 (62249, Thermo Fisher Scientific), and coverslips were mounted using ProLong Gold Antifade reagent (P36930, Thermo Fisher Scientific). Imaging was conducted with a Leica STELLARIS 8 FALCON microscope (Leica Microsystems, Wetzlar, Germany) equipped with an HC PL APO 100x/1.40 oil CS2 objective (11506372, Leica Microsystems). Excitation light intensity was set to 2% for Hoechst (λ_ex_ = 405 nm; λ_em_ = 425-622 nm), 3% for Alexa488 (λ_ex_= 499 nm; λ_em_ = 504-587 nm), and 50% for NPY-mCherry (λ_ex_ =587 nm; λ_em_= 593-750 nm). Typical exposure was 200 ms. Images were acquired using Leica Application Suite X (LAS X), and analysis was performed with ImageJ software^178^. Co-localization analysis was conducted using the JACoP plugin^179^ in ImageJ.

### Bulk release of serotonin

Upon reaching ∼80% confluency, BON cells (passages 2-10), were trypsinized and plated in 24-well cell culture plates pre-coated with poly-L-lysine followed by 15 µg/ml of laminin (Sigma-Aldrich, Burlington, MA). ∼150,000-200,000 cells were added in each well and allowed to attach to the plate surface. 300 µM of serotonin (Sigma) was added to the each well and incubated for 24-48 h at 37°C under 5% CO_2_. After 48 hours of incubation, DMEM/F12 medium was removed, and the cells were thoroughly washed 3-5 times with pre-warmed PBS followed by one wash with pre-warmed amperometry buffer (AB, 140 NaCl, 5 KCl, 5 CaCl2, 1 MgCl2, 10 HEPES/NaOH, pH 7.3). For each experiment two parallel sets of wells were selected. In one set of wells, 200 μl of ionomycin (10 μM) in AB was added and incubated for ∼3 s. The supernatant was immediately collected without touching the cells and centrifuged at 15000 g for 1 hr at 4°C. An aliquot of 50 μL of the supernatant was collected and used for the serotonin ELISA assay. In another set of wells, cells were trypsinized and collected as a pellet and resuspended in 500 μL of resuspension buffer (150 KCl, 20 HEPES, 20 EDTA, EDTA free protease inhibitor cocktail). The cells were passed through a 27 gauge needle at least 20 times and centrifuged at 15000 g at 4°C for 2 h. The supernatant fraction was collected and a 50 µl aliquot (total cell lysate) of the supernatant was used for the serotonin ELISA assay.

Estimation of serotonin concentration in the supernatant was done using an Invitrogen serotonin competitive ELISA kit (EEL006, Thermo Fisher Scientific). The assay was carried out according to the manufacturer’s protocol and the concentration of the serotonin was estimated against the serotonin standards carried out in parallel. The final concentration of serotonin was adjusted based on the dilution and percent secretion was calculated as a ratio of serotonin in the Ionomycin induced to that of the total cell lysate. All experiments were performed in dark to avoid exposure to light as much as possible. The experiment was repeated 3 times.

### Amperometry

Thirty-five mm diameter glass-bottom dishes (with 14 mm diameter, #1.5 glass, poly-D-lysine coated, P35GC-1.5-14-C, Mattek, Ashland, MA) were further coated with laminin (15 µg/ml, Sigma) by incubation for 1 hour. After incubation, excess laminin was removed, and the dish was gently washed with PBS. followed by a gentle wash with PBS. About 10.BON cells were plated on this dish and incubated for couple of hours for cell attachment. Serotonin (300 µM) was then added to the medium and incubated for 24-36 hours. For amperometry recordings, cells were washed 2-3 times with AB and then 3 ml of AB was added to the cells and the dish was transferred to the microscope stage for amperometry experiments.

An Olympus IX71 microscope (Olympus America, Waltham, MA) equipped with a water immersion objective (UApoN340, 40x 1.15W) and a Prior Lumen 200 Pro excitation light source was used to locate cells, place the carbon fiber electrode (CFE) and the puffing pipette. NPY-mCherry fluorescence was used to locate granules within cells with excitation and emission filters transmitting 542-582 nm and 600-652 nm, respectively.

A freshly cut carbon fiber electrode (CFE, 5 µm diameter, ALA Scientific, Farmingdale, NY) was placed near a cell, gently touching it, using an MP-285 micromanipulator controlled by an MPC-200 controller (Sutter Instruments, Novato, CA). The pipettes for delivering the stimulation solution were pulled from 1.5 mm outside diameter borosilicate glass capillaries (BF150-86-10, Sutter Instruments) using a P-1000 pipette puller (Sutter Instruments) and had 1-2 µm diameter openings. Ionomycin stock solution (1 mM) was diluted to working concentration of 10 µM in AB and was filled in the puffing pipettes devoid of any air bubbles. The puffing pipette was placed approximately 10 µm from the cell. An oxidizing potential of +650 mV (vs Ag/AgCl) was continuously applied to the CFE via a HEKA EPC10 USB amplifier controlled *via* Patchmaster software (HEKA Electronic, Harvard Apparatus, Holliston, MA). Pressure-driven superfusion of the stimulation solution was achieved using a Picospritzer III instrument (Parker Hannifin Co., Hollis, NH). The timing of stimulation was controlled by sending a voltage signal from the HEKA EPC10 to Picospritzer III. Ionomycin was applied for 3 s after recording of a baseline for 6 s. Recording continued for 14-25 s after stimulation ended. Currents were recorded with gain = 50 mV/pA, sampled at 20 kHz and filtered by a 10 kHz and 3 kHz Bessel filters, typically resulting in ∼ 1 pA root-mean squared noise.

Amperometric traces were exported from Patchmaster as Igor Pro files (Wavemetrics, Portland, OR) and analyzed using Igor Pro procedures written by Mosharov and Sulzer^68^, using the default settings. Statistics of individual spike parameters were further analyzed using MATLAB. Parameters were first averaged for a given cell, then averaged across cells^180^. Fusion pore permeability was calculated following Jackson et al.^75^

### Calcium measurements

We used the parent, unlabeled N13 clone under identical conditions as the cells co-expressing pH-Syt1 constructs and NPY-mCherry to avoid overlap between fluorescence signals. Cells were incubated with 5 µM Fluo-4-AM (Thermo Fisher scientific, Waltham, MA) along with 0.1% pluronic F-127 (Thermo Fisher Scientific) in HBSS buffer for 1 hr, followed by a gentle wash with PBS and incubation in DMEM/F12 medium for 30 min. Cells were then transferred to the microscope described above, equipped with a photomultiplier tube (PMT) detection system (model 810, Photon Technology International, Horiba Scientific, Piscataway, NJ). Fluo-4 fluorescence was excited and collected through a GFP set (472/30 excitation filter and 544/47 emission filter from Semrock Brightline GFP-3035B-OMF filter set, Semrock Optical filters, IDEX Health & Science LLC., Rochester, New York) and detected using the PMT. Signals were collected from a limited area that included a single cell, selected using a diaphragm. Similarly, excitation was limited only to an area slightly larger than the cell under examination.

Background signals were recorded from unlabeled, parental BON N13 cells that were not loaded with dye using otherwise identical conditions. The average background level was subtracted from the fluorescence traces. Then changes in the fluorescence relative to the initial signal *F* −*F*_0_ = Δ*F*/*F*_0_ were calculated, where *F*_0_ is the initial fluorescence level before stimulation.

### Electron microscopy

BON cells were seeded on laminin-coated coverslips, and then fixed with 5% glutaraldehyde in 0.1 M sodium cacodylate buffer which was added directly to the media. The solution was replaced with 2.5% glutaraldehyde in 0.1 M sodium cacodylate buffer (pH 7.4), and the samples were washed with cacodylate buffer. Next, the samples were incubated with 0.5% osmium tetroxide, 0.8% potassium ferrocyanide, rinsed with 0.1 M sodium cacodylate and HPLC water before undergoing en bloc staining in 2% uranyl acetate in 0.1 M sodium cacodylate. The samples underwent ethanol dehydration in steps of 50%, 70%, 90%, 3 x 100% anhydrous ethanol 1:1 (v/v), followed by 100% ethanol in EPON. The samples were then embedded in 100% EPON epoxy resin and polymerized overnight at 60°C. The samples were sectioned into 60 nm thick sections using a Leica EM7 Ultramicrotome and placed on formvar-coated nickel mesh grids. The grids were post-stained with 2% uranyl acetate and Reynolds Lead citrate, and imaged on a Tecnai 12 Biotwin Transmission EM at 80 kV using an AMT camera. For stimulated cells, the same procedure was used, except 1 µM ionomycin was applied for ∼3 s just before addition of the fixative.

Electron micrographs were analyzed using Image J^178^. To quantify areas, DCV outlines were traced using the magic wand tool, or the polygon selection tool followed by spline fitting. For measuring the minimum distance *d* between a DCV and the PM, we used the straight line tool and drew the closest distance between the two membranes. Results were saved in a spreadsheet for further analysis. At least two independent cell cultures and sample preparations were used for each condition. For the distribution of DCV-PM distances, we only considered distances ≤300 nm, roughly equal to one DCV diameter. Note that the distances *d* we measured cannot be taken literally, as they are affected by chemical fixation and sectioning artifacts. However, these artifacts should affect all the samples equally, so *d* can be used as a qualitative measure of differences among groups. A careful quantification of *d* requires specialized equipment that allows rapid freezing after stimulation.

### Statistical analyses

Unless otherwise noted, we used the non-parametric Kruskal-Wallis test to compare the medians of the groups of data to determine if the samples came from the same distribution. Unlike 1-way ANOVA, this test uses ranks of the data to compute the test statistics, and does not assume the data follow normal distributions. To determine which groups of data had different mean ranks compared to the control group (usually the parental BON N13 cells, or WT rescues) we used the multiple comparison test using Dunnett’s procedure, or to determine differences among groups, the Tukey-Kramer procedure.

## ACKNOWLEDGEMENTS

We thank the members of the Karatekin and Toomre labs for valuable discussions, and Bruno Gasnier for the BON N13 clone. We are grateful to D. Zenisek (Yale University) for discussions and feedback on the manuscript. We acknowledge funding from the National Institutes of Health, National Institute of Neurological Disorders and Stroke (grant R01 NS113236 to EK). The funders had no influence in the design, execution, and interpretation of the study.

## AUTHOR CONTRIBUTIONS

MT, DT, FRM, and EK conceived the study. FRM generated the new BON cell lines and characterized them for expression. MT and SKC performed the amperometry experiments and the associated data analysis. DC performed EM experiments and analysis. AL performed colocalization studies. MT, SKC, DC, FRM, AL and EK analyzed data. EK and DT acquired funding and supervised the work. EK wrote the manuscript with input from all authors.

